# Phylogenomic Taxonomic Analysis of *Ralstonia solanacearum* Strains causing Bacterial Wilt Disease in Northeastern Argentina

**DOI:** 10.64898/2026.04.29.721750

**Authors:** Veronica Obregon, Gi Yoon Shin, Ernestina Galdeano, Romina Escobar, Tatiana Lattar, Julia Magali Ibañez, Ariel Amadio, Jose Matias Irazoqui, Gonzalo Manuel Santiago, Maria Florencia Eberhardt, Alberto Martin Gochez, Tiffany M. Lowe-Power

## Abstract

*Ralstonia solanacearum* species complex (RSSC) is a genetically diverse group of plant pathogens, yet genomic data from South America remain limited. Here, we characterize 13 RSSC strains isolated from tomato, pepper, and eggplant in northeastern Argentina.

Phylogenetic analysis of the *egl* marker gene assigned these strains to phylotype IIA and suggested two closely related lineages. Complete genomes (5.63–5.76 Mb) were generated for four representative strains, yielding high-quality (99.94% completeness with f_Burkholderiaceae CheckM markers), closed assemblies with canonical bipartite architecture. Phylogenetic analysis of the *egl* marker, 49 conserved bacterial genes, and average nucleotide identity (ANI) analyses, consistently assigned one lineage to sequevar IIA-50, forming a coherent and monophyletic group. In contrast, although *egl* analysis suggested the second lineage was related to one sequevar IIA-38 reference strain, genomic analysis did not support this assignment. Further, the genomic analysis revealed significant genomic distance between the genomes for two sequevar 38 representative strains, supporting a conclusion that sequevar 38 itself was not monophyletic and instead appears paraphyletic. These findings highlight limitations of single-locus classification and support genome-informed refinement of RSSC sub-phylotype taxonomy.

**Outcome statement:** Reports of bacterial wilt disease in Argentina had not yet been published in the international literature although the disease has been long-standing. This study provides complete genome sequences for four *Ralstonia solanacearum* strains from Northern Argentina and places them within a global phylogenomic framework. The Argentine strains cluster into two closely related phylotype IIA lineages, indicating that bacterial wilt in this regional dataset is associated with genetically similar populations. For clear communication of which strains are present in Northern Argentina, we attempted to classify the lineages to the long-standing sequence variant (sequevar) system for naming *R. solanacearum* species complex (RSSC) strains. One lineage was confidently assigned to IIA-50 with genomic support that confirmed phylogenetic analysis of the classical genetic marker *egl*. However, newly available genomes for sequevar reference strains revealed an issue where two distantly related strains are currently recognized as references for sequevars. Overall, these results provide evidence supporting the need for genome-informed refinement of sub-phylotype classification and expand genomic representation of South American RSSC populations.

**Data summary:** Complete genome assemblies and raw reads for INTABV18, INTABV29, INTABV624 and INTABV2657 are deposited to NCBI under the project number PRJNA1407867. The curated dataset of public RSSC genomes is available to users who register a free account on KBase via a KBase narrative (https://narrative.kbase.us/narrative/189849). The narrative described in a living BioRxiv pre-print [1]. Supplemental files such as Figure S1, rectangular versions of all trees (Figure 2 and 3 and S1) and supplementary table S1, S2, S3 and S4 are available on Zenodo at doi.org/10.5281/zenodo.19502890

## Introduction

Bacterial plant pathogens in the *Ralstonia solanacearum* species complex (RSSC) invade the xylem vasculature, leading to wilt disease. The RSSC is genetically diverse and composed of three species and four phylotypes: *R. solanacearum* (phylotype II), *R. pseudosolanacearum* (phylotype I and III), and *R. syzygii* (phylotype IV) [2–5]. Although there is no perfect classification system for identification and naming of RSSC lineages, the sequevar (sequence variant) system remains valuable because it uses DNA sequence comparisons, has been widely applied to RSSC strains worldwide, and there is a recently published standardized procedure and reference sequences [6]. Sequevars are determined through phylogenetic comparison using a specific portion of the *egl* endoglucanase gene, a conserved pathogenicity gene [2, 7, 8]. Although there are documented cases of phylogenetic incongruence between sequevar/*egl*-typing and genomic analysis [9, 10], sequevar identification remains useful for pathogen surveillance due to the lower cost relative to whole genome sequencing.

Collectively, the RSSC has a global distribution with strain isolations reported in over 107 countries and territories [1]. Although the phylotypes have originated in different locations (phylotype I, Asia; phylotype II, the Americas; phylotype III, Africa; and phylotype IV, Indonesia, Australia, and Oceania) [2], commercial trade of plant parts has led to the dispersal and establishment of certain lineages in new countries and continents. Currently, phylotypes I and II have a broad geographic distribution with presence on all continents with plant agriculture, phylotype III remains limited to Africa, and phylotype IV is predominately limited to Oceania and Asia except for recent reports in Kenya and Brazil [11–14].

In Argentina, the RSSC has been reported in multiple provinces on banana, plantain, tomato, potato, and tobacco in regional journals that are not indexed in major international databases (Figure 1). RSSC pathogens were first reported in Argentina in 1919, where epidemics of banana and plantain wilt were described in the Northeastern provinces of Chaco, Formosa, Corrientes, and Misiones [15]. Bacterial wilt of tomato was first reported in 1972 in Corrientes [16]. In the 1990s, this disease was reported in the provinces of Santa Fe, Formosa, Entre Ríos, Santa Fe, Misiones, and Tucumán [17–19]. During 2005, bacterial wilt of tomato was reported in the provinces of Jujuy and Salta [20]. RSSC pathogens are reported on tobacco in the provinces of Jujuy, Salta, and Misiones [20–22]. Brown rot of potato was first reported during the 1978-1979 season in southeast of the Buenos Aires province [23]. The movement of seed potatoes between different production areas in Argentina and abroad likely led to the unintentional introduction of this new disease. The potato brown rot pathogen in Argentina was identified as race 3 biovar 2 [23], which is modernly recognized as *R. solanacearum* phylotype IIB sequevar 1 or sequevar 2 [2, 24].

**Figure 1.**
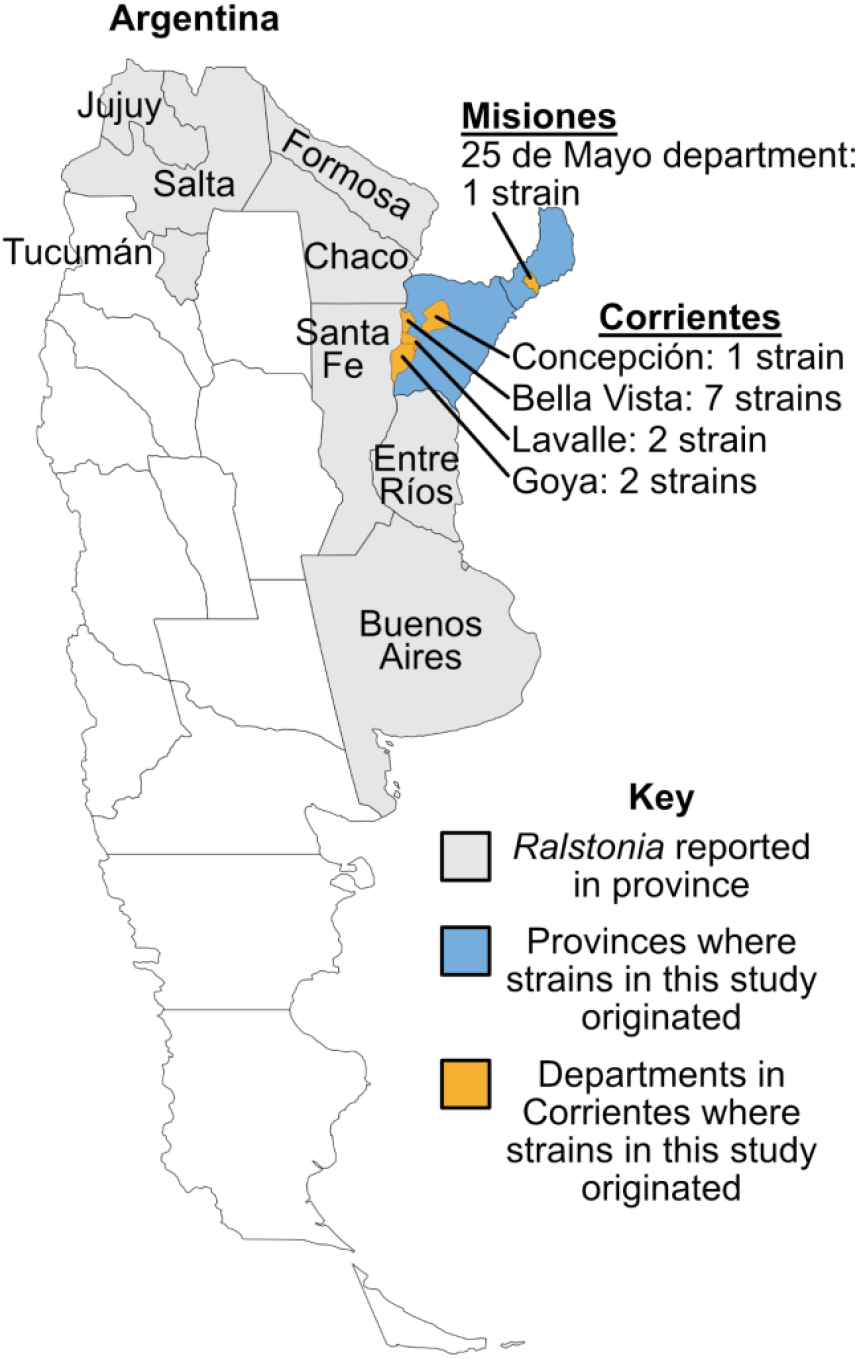
Map of locations where RSSC wilt disease has been documented in this study and in the literature. Provinces where RSSC incidence has been reported are labeled: Jujuy, Salta, Tucumán, Formosa, Chaco, Santa Fe, Buenos Aires, Misiones, Corrientes, and Entre Ríos. Strains analyzed in this study were collected from two provinces (blue): Corrientes (departments of Concepción, Bella Vista, Lavalle, and Goya) and Misiones (department of 25 de Mayo), both indicated in yellow.

**Figure S1.**
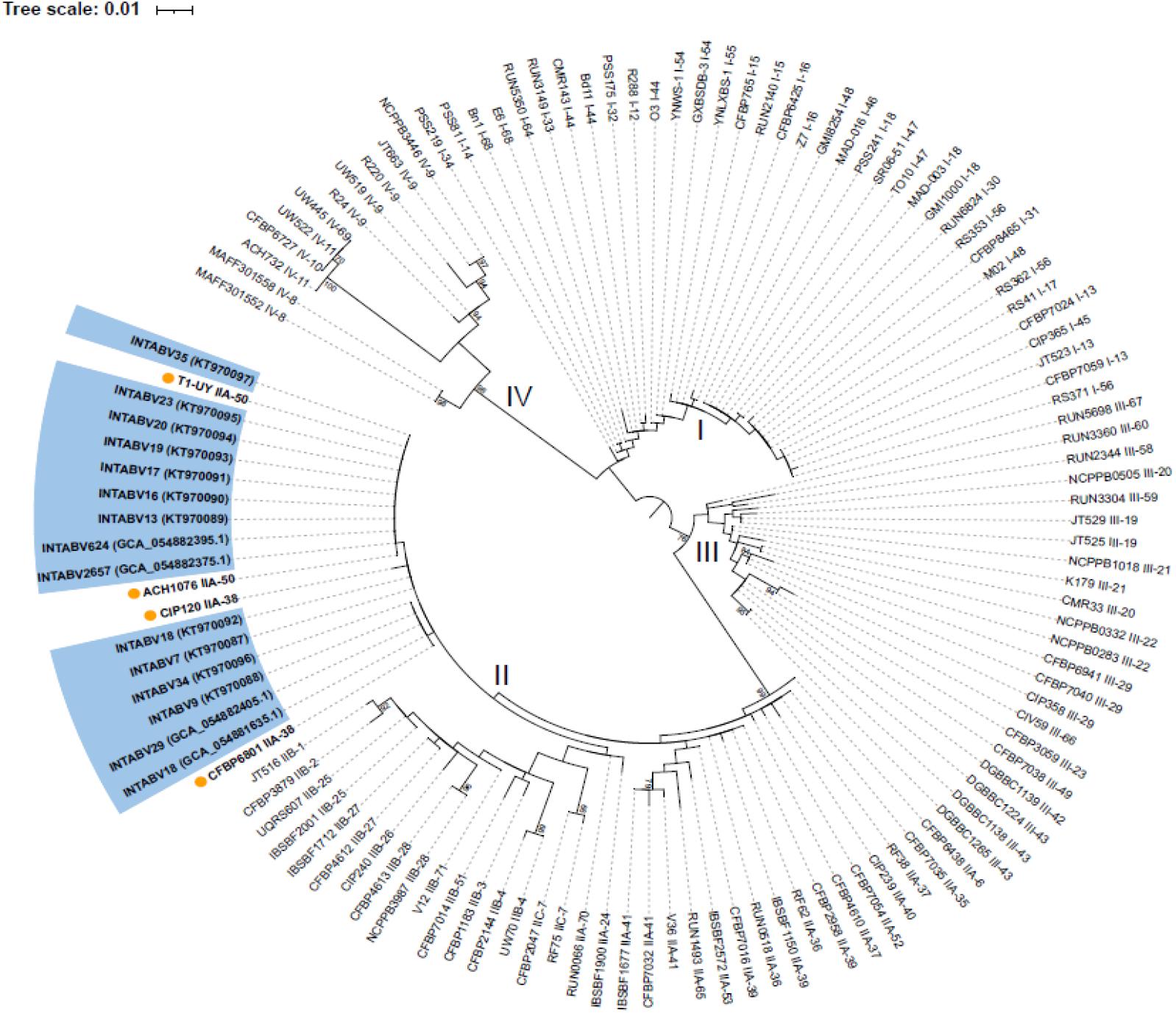
Maximum-likelihood phylogenetic tree inferred from 471 bp of endoglucanase (*egl*) gene sequences assigned Argentine strains as phylotype II sequevar 38 and sequevar 50. The tree was constructed using PhyML v3.0 under the GTR nucleotide substitution model with gamma-distributed rate heterogeneity (α = 0.33), as selected by the SMART model selection procedure implemented in PhyML (Lefort *et al*., 2017). The *egl* sequences from Argentine strains are highlighted in blue, and their corresponding GenBank accession numbers for both the *egl* nucleotide sequence and the whole-genome assembly are shown in parentheses. Reference *egl* sequences representing sequevars IIA-38 (CFBP6801 and CIP120) and IIA-50 (T1-UY and ACH1076) are also shown in bold and marked with yellow circles. A searchable PDF of this tree in rectangular format is available on Zenodo (doi.org/10.5281/zenodo.19502890).

## Methods

### Strain isolation

Commercial greenhouse and field crops were surveyed in Bella Vista, Colonia 3 de Abril, Colonia Progreso, Santa Lucía, Lavalle, Goya, Empedrado, Saladas, Tata Cuá, Gobernador Martínez, and other localities within the province of Corrientes in Argentina. These strains were isolated at the Horticultural Plant Pathology Laboratory of the INTA Bella Vista Agricultural Experimental Station during 2008 (INTABV29; INTABV18), 2012 (INTABV624), and 2024 (INTABV2657).

The stems and roots of eggplant, pepper and tomato plants displaying wilting symptoms were sampled for isolation. Plant materials were first thoroughly rinsed under running tap water, then surface-disinfected in 0.5% sodium hypochlorite for 1 min, followed by three consecutive rinses in sterile distilled water. A second disinfection step was performed using 70% ethanol for 1 min followed by three rinses in sterile water to remove residual disinfectants. Disinfected stem and root tissues were cut into small fragments and placed in test tubes containing sterile distilled water to promote bacterial streaming [25]. When plant material was not fresh, tissue fragments were macerated in a mortar with sterile distilled water prior to plating. Aliquots were streaked onto CPG-TZC (hydrolyzed casein (1 g/L), peptone (10 g/L), glucose (5 g/L), agar (17 g/L), and tetramethyl-triphenyl-tetrazolium (0.005%) [26] medium using a 5 μL inoculating loop. Inoculated plates were incubated at 28 °C for 2–3 days.

### Genomic DNA Extraction, Assembly and Strain Selection

Genomic DNA was extracted using the CTAB method as described by [27]. The four sequenced strains were isolated from tomato (Corrientes province: strain INTABV29, from Lavalle in 2008; strain INTABV624 from 3 de Abril in 2012; strain INTABV2657 from Yatayti Calle in 2024) and pepper (Misiones province: strain INTABV18 in 2008). Strains were selected to include different hosts, geographic locations, and years of isolation within *R. solanacearum* phylotype IIA populations previously identified in the region. They were whole genome sequenced using long-read Oxford Nanopore Technologies (ONT). High molecular weight (HMW) genomic DNA was end-repaired, and libraries were constructed using Native Barcoding Kit v14 (SQK-NBD114.24, ONT) following the manufacturer’s protocol. Libraries were loaded onto an R10.4.1 flow cell (FLO-MIN114) and sequenced on a MinION device. Basecalling was performed using dorado v1.1.1 (github.com/nanoporetech/dorado) with the high-accuracy super (SUP) model v5.2.0.

Reads were filtered using Filtlong v0.3.1 (github.com/rrwick/Filtlong) and then assembled *de novo* with Flye v2.9.6-b1802, applying default settings [28]. The genome assemblies were annotated with NCBI Prokaryotic Genome Annotation Pipeline (PGAP) [29] while their quality and completeness were assessed using KBase version QUAST v4.4 [30, 31] and CheckM v1.0.18 with the markerset for the Burkholderiaceae family [32].

### PCR and Sanger Sequencing of Partial *egl* Sequence

Partial *egl* gene fragments were amplified using the EndoF/EndoR primer pair as described by [33]. PCR reactions were carried out in a final volume of 25 µl containing 1× reaction buffer (200 mM Tris-HCl, pH 8.4; 500 mM KCl), 0.75 mM MgCl_2_, 0.2 mM of each dNTP, 0.2 µM of each primer, 1 U of Taq DNA polymerase, and 100–150 ng of template DNA. Amplifications were performed with an initial denaturation at 95 °C for 5 min, followed by 30 cycles of 95 °C for 1 min, 67 °C for 1 min, and 72 °C for 2 min, with a final extension at 72 °C for 10 min. PCR products were purified, end-repaired using Klenow fragment (Promega), and cloned into the pBluescript II SK+ vector (Stratagene) digested with Sma*I*. Recombinant plasmids were transformed into *Escherichia coli* DH5α and extracted by alkaline lysis. Inserts were sequenced using M13 universal primers at the Genomics Unit (Instituto de Biotecnología, CICVyA, CNIA, INTA, Argentina).

### Phylogenetic analyses

RSSC sequevar assignment was performed by comparing partial sequences of the endoglucanase (*egl*) gene between the strains in this study and a curated reference dataset of 107 partial *egl* sequences [2, 6]. A recently published standard operating procedure recommends that *egl* comparisons use the gold-standard 678 to 710 bp region of the gene [7], which we were able to analyze for the four strains with genome sequences. We also had existing, unpublished Sanger sequencing data for one of the genome-sequence strains (INTABV18) and ten additional strains with which we conducted a less precise *egl* analysis of a shorter 471 bp region. The partial *egl* sequences were aligned using MAFFT implemented in Geneious Prime v2026.02 and the alignment was trimmed to a final length of 471 basepairs (bp) or to the locations recommended in [7]. The trimmed alignment was exported in PHYLIP format and subjected to phylogenetic analysis using online PhyML v3.0 [34]. The best-fit nucleotide substitution model was selected according to the Akaike Information Criterion (AIC). A maximum-likelihood phylogeny was inferred with 1,000 bootstrap replicates to assess branch support. The resulting *egl* gene tree was visualized and annotated using iTOL v7.4.2 [35].

To assess the phylogenetic relatedness of the four INTABV genomes sequenced in this study relative to global RSSC diversity, we compared them to a dataset of 826 public RSSC genomes available through an open access KBase narrative for RSSC Phylogenomics that we maintain [1]. This public data includes the *egl* reference strains that were useful in this study: CIP120 from [24], CFBP6801 recently sequenced in [36], and T1-UY originally isolated by María Inés Siri (personal communication) and sequenced in [24]. Phylogenetic reconstruction was performed using the Insert Genomes Into a SpeciesTree v2.2.0 in KBase, which uses FastTree2 to infer an approximate maximum-likelihood tree based on 49 conserved single-copy marker genes [31, 37]. The set of marker genes used by the SpeciesTree pipeline is publicly documented within the KBase application framework https://kbase.us/applist/apps/SpeciesTreeBuilder/insert_genomeset_into_species_tree/release) and listed in [38]. The resulting species tree was exported and visualized using iTOL.

To quantify genomic relatedness between genomes of the Argentine strains and closely related genomes, and phylotype reference genomes, 41additional publicly available assemblies were included for Average Nucleotide Identity (ANI) analysis (listed in Table S1). Pairwise ANI values were calculated using FastANI v1.33 [39] using default parameters and a heat map was visualized using Conditional Formatting in Microsoft Excel.

## Results

### Genome statistics for the four complete, Argentine *R. solanacearum* genomes

We selected four RSSC strains from Northern Argentina for whole-genome sequencing and analysis. Nanopore sequencing and assembly of INTABV624, INTABV29, INTABV18, and INTABV2657 yielded high-quality, complete genomes with uniform sequencing depth (~52× coverage) (Table 2). Each genome assembled into two circular replicons corresponding to the chromosome and the megaplasmid, consistent with the canonical genome architecture of *Ralstonia* species [40]. Genome sizes ranged from 5.63 Mb (INTABV624) to 5.76 Mb (INTABV2657). The largest replicons and corresponding N50 values represent the chromosomal component: 3.47–3.49 Mb. GC content was highly consistent across strains (66.32–66.50 %). Gene prediction with NCBI PGAP identified 5,049–5,130 total genes per genome, including 4,987–5,062 coding sequences (CDS) and 50–122 putative pseudogenes. The number of predicted protein-coding genes ranged from 4,865 to 4,981 and ribosomal RNA gene counts were similar across strains (66–68 rRNA genes). Genome quality assessment using CheckM with the f Burkholderiaceae marker sets indicated the assemblies were high quality assemblies with 99.94% completeness and minimal contamination (0.46%), supporting the integrity and completeness of both chromosomal and megaplasmid sequences.

### *R. solanacearum* strains infecting tomato, pepper and eggplant in Northeastern Argentina belong to the monophyletic IIA-50 sequevar or a closely related lineage

We investigated phylogenetic relationships of the Argentine strains via sequence analysis of the standard RSSC marker gene (*egl*) and genomic approaches. As described below, all analyses consistently showed that there are two closely related phylotype IIA lineages in Northeastern Argentina. However, the *egl* and genomic analyses did not agree on the relationships of the Argentine strains with certain sequevar reference strains.

We investigated traditional sequevar assignments for the four genomes, using the 710 bp region of *egl* recommended in a recently published standard operating procedure [7]. A maximum-likelihood tree clustered two Argentine strains with IIA-50 reference strains T1-UY (Uruguay) and ACH1076 (Brazil): INTABV624 and INTABV2657 (Figure 1). The other two Argentine strains (INTABV18 and INTABV29) clustered with one IIA-38 reference strain (CFBP6801 from Martinique Island) and two non-reference IIA-38 strains from the Southeastern USA (Rs124 and UCD576). Unexpectedly, the second IIA-38 reference strain (CIP120 from Peru) was not monophyletic with this IIA-38 cluster. Hereafter, we tentatively refer to the Argentine lineages as IIA-50 and “clade 2” due to the uncertainty of whether IIA-38 is a monophyletic lineage. In previous years, we had performed Sanger sequencing of the *egl* gene on 14 Argentine strains and the sequence was trimmed to a 471 bp region that was high-quality in all the Sanger traces, which is shorter than the recommended length [7]. By this analysis, the additional Argentine strains had clustered with the genome-sequenced Argentine IIA-50 or “clade 2” strains (Figure S1), so we tentatively assigned these either IIA-50 or “clade 2” (Table 1). These assignments should be treated with caution because 471 bp provides low discriminatory power, as demonstrated by the Figure S1 tree failing to accurately resolve the well-documented separation of the phylotype IIA and IIB lineages.

**Table 1.**
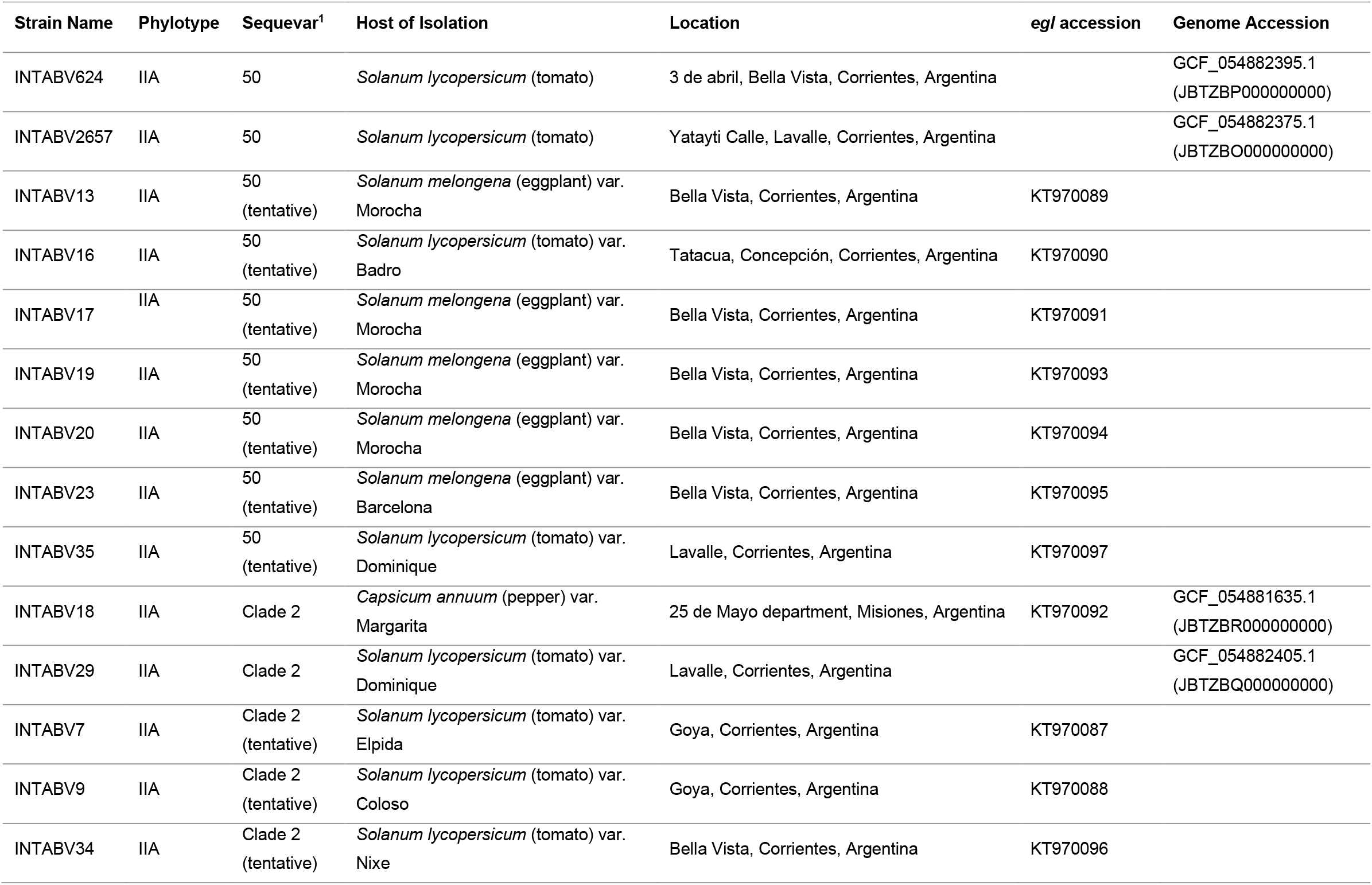
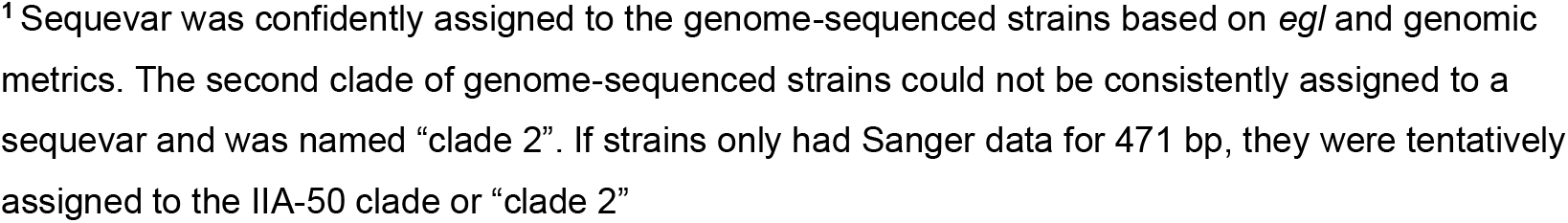
Strains used in this study.

| Strain Name | Phylotype | Sequevar<sup>1</sup> | Host of Isolation | Location | egl accession | Genome Accession |
| --- | --- | --- | --- | --- | --- | --- |
| INTABV624 | IIA | 50 | *Solanum lycopersicum* (tomato) | 3 de abril, Bella Vista, Corrientes, Argentina |  | GCF_054882395.1<br>(JBTZBP000000000) |
| INTABV2657 | IIA | 50 | *Solanum lycopersicum* (tomato) | Yatayti Calle, Lavalle, Corrientes, Argentina |  | GCF_054882375.1<br>(JBTZBO000000000) |
| INTABV13 | IIA | 50<br>(tentative) | *Solanum melongena* (eggplant) var.<br>Morocha | Bella Vista, Corrientes, Argentina | KT970089 |  |
| INTABV16 | IIA | 50<br>(tentative) | *Solanum lycopersicum* (tomato) var.<br>Badro | Tatacua, Concepción, Corrientes, Argentina | KT970090 |  |
| INTABV17 | IIA | 50<br>(tentative) | *Solanum melongena* (eggplant) var.<br>Morocha | Bella Vista, Corrientes, Argentina | KT970091 |  |
| INTABV19 | IIA | 50<br>(tentative) | *Solanum melongena* (eggplant) var.<br>Morocha | Bella Vista, Corrientes, Argentina | KT970093 |  |
| INTABV20 | IIA | 50<br>(tentative) | *Solanum melongena* (eggplant) var.<br>Morocha | Bella Vista, Corrientes, Argentina | KT970094 |  |
| INTABV23 | IIA | 50<br>(tentative) | *Solanum melongena* (eggplant) var.<br>Barcelona | Bella Vista, Corrientes, Argentina | KT970095 |  |
| INTABV35 | IIA | 50<br>(tentative) | *Solanum lycopersicum* (tomato) var.<br>Dominique | Lavalle, Corrientes, Argentina | KT970097 |  |
| INTABV18 | IIA | Clade 2 | *Capsicum annuum* (pepper) var.<br>Margarita | 25 de Mayo department, Misiones, Argentina | KT970092 | GCF_054881635.1<br>(JBTZBR000000000) |
| INTABV29 | IIA | Clade 2 | *Solanum lycopersicum* (tomato) var.<br>Dominique | Lavalle, Corrientes, Argentina |  | GCF_054882405.1<br>(JBTZBQ000000000) |
| INTABV7 | IIA | Clade 2<br>(tentative) | *Solanum lycopersicum* (tomato) var.<br>Elpida | Goya, Corrientes, Argentina | KT970087 |  |
| INTABV9 | IIA | Clade 2<br>(tentative) | *Solanum lycopersicum* (tomato) var.<br>Coloso | Goya, Corrientes, Argentina | KT970088 |  |
| INTABV34 | IIA | Clade 2<br>(tentative) | *Solanum lycopersicum* (tomato) var.<br>Nixe | Bella Vista, Corrientes, Argentina | KT970096 |  |

<sup>1</sup> Sequevar was confidently assigned to the genome-sequenced strains based on *egI* and genomic metrics. The second clade of genome-sequenced strains could not be consistently assigned to a sequevar and was named “clade 2”. If strains only had Sanger data for 471 bp, they were tentatively assigned to the IIA-50 clade or “clade 2”

**Table 2.**
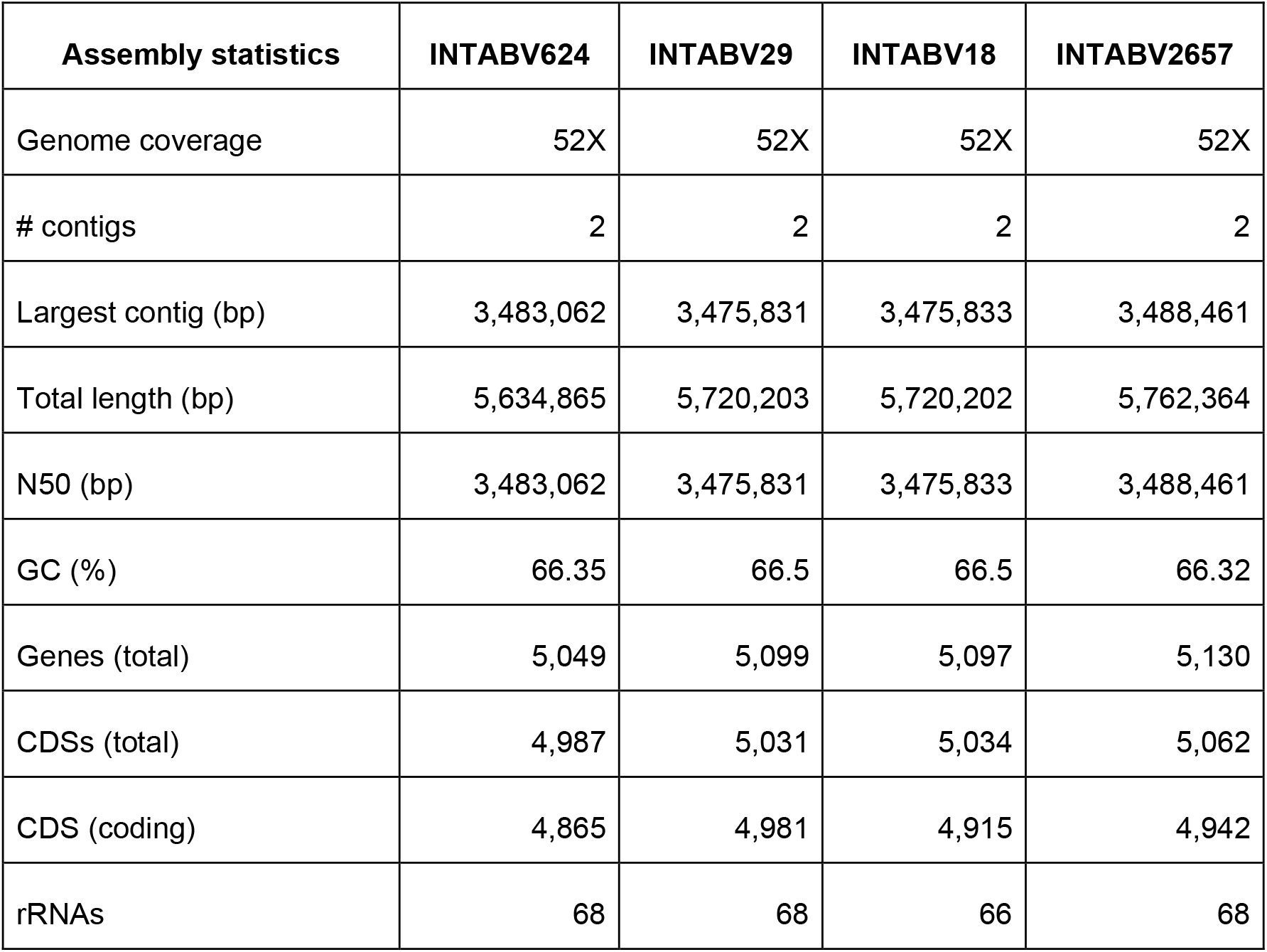
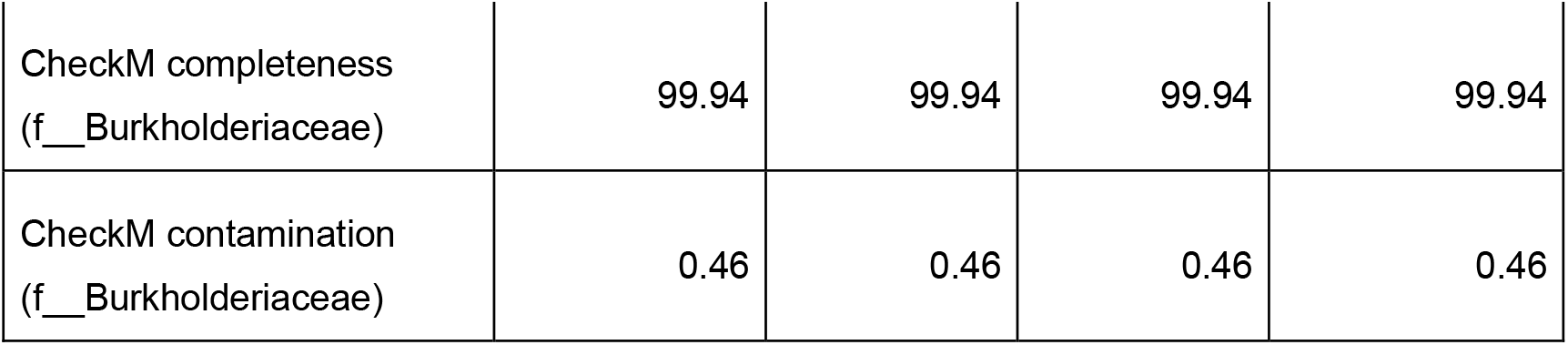
Genome assembly statistics for complete genomes of four *R. solanacearum* strains from Argentina.

| Assembly statistics | INTABV624 | INTABV29 | INTABV18 | INTABV2657 |
| --- | --- | --- | --- | --- |
| Genome coverage | 52X | 52X | 52X | 52X |
| # contigs | 2 | 2 | 2 | 2 |
| Largest contig (bp) | 3,483,062 | 3,475,831 | 3,475,833 | 3,488,461 |
| Total length (bp) | 5,634,865 | 5,720,203 | 5,720,202 | 5,762,364 |
| N50 (bp) | 3,483,062 | 3,475,831 | 3,475,833 | 3,488,461 |
| GC (%) | 66.35 | 66.5 | 66.5 | 66.32 |
| Genes (total) | 5,049 | 5,099 | 5,097 | 5,130 |
| CDSs (total) | 4,987 | 5,031 | 5,034 | 5,062 |
| CDS (coding) | 4,865 | 4,981 | 4,915 | 4,942 |
| rRNAs | 68 | 68 | 66 | 68 |
| CheckM completeness<br>(f__Burkholderiaceae) | 99.94 | 99.94 | 99.94 | 99.94 |
| CheckM contamination<br>(f__Burkholderiaceae) | 0.46 | 0.46 | 0.46 | 0.46 |

We placed the Argentine strains’ genomes into a phylogenetic context with 826 public genomes from global RSSC strains using the KBase SpeciesTree pipeline that analyzes 49 conserved genes. The boxed clade in Figure 3 comprises genomes of the IIA sub-phylotype, including some genomes with sequevar assignments. For the IIA-50 strains, the SpeciesTree was congruent with the *egl* analysis; INTABV624 and INTABV265 grouped with genomes of IIA-50 reference strain TI-UY from Uruguay, seven Brazilian strains, and one strain isolated from an unknown location. The Argentine clade 2 formed a distinct branch that was sister to the IIA-50 clade although bootstrap support was moderate (0.731; Figure 3). The Southeastern USA IIA-38 strains were clustered near the Argentine strains. In contrast, the genomes of IIA-38 reference strains CIP120 and CFBP6801 were dispersed across the IIA branch.

Due to the irregularities between the *egl* tree and the 49-gene tree, we further assessed genome-wide relatedness by calculating pairwise Average Nucleotide Identity (ANI) for the four Argentine strains with 41 pubic IIA genomes (Figure 4). Once again, this analysis was simpler to interpret for the IIA-50 sequevar than for the non-monophyletic IIA-38 sequevar. INTABV624 and INTABV2657 shared very high ANI values (>99.94%) with the nine tested IIA-50 genomes, confirming their assignment to sequevar IIA-50 by this third method (Figure 4). Clade 2 strains INTABV18 and INTABV29 had nearly identical ANI with each other (99.99955%). Similar to how INTABV18 and INTABV29 branched closely to the monophyletic IIA-50 clade on the 49-gene tree, ANI between these two strains and the eleven IIA-50 strains was relatively high at ~99.52%. In contrast, the IIA-38 reference strains had considerable genetic distance to any Argentine strain at ~98.79% ANI to the clade 2 genomes (INTABV18 and INTABV29) and 98.81% ANI to the IIA-50 genomes (INTABV624 and INTABV2657). Ultimately, we decided to refrain from placing the clade 2 Argentine strains into a named sequevar.

## Discussion

This study is the first to phylogenomically investigate RSSC strains from Argentina.

Fourteen strains isolated from eggplant, pepper, and tomato in Corrientes (n = 13) and Misiones (n = 1) provinces in Argentina were genotypically identified as *R. solanacearum* strains belonging to two closely related clades of phylotype IIA. Although we set out with the simple goal of identifying the sequevar of both lineages, we achieved a confident taxonomic assignment for only the IIA-50 lineage. For the scope of this report, we use “clade 2” as a placeholder for this second lineage.

The development of the phylotype and sequevar system has improved the ability of scientists to communicate about RSSC diversity with reasonable accuracy [2, 11, 24, 41, 42]. Nevertheless, sequevars are based on less than 1 kb of sequence of a single marker gene, and even the original definition addressed the limitations: “a sequevar, or sequence variant, is defined as a group of strains with a highly conserved sequence within the area sequenced.” The discrepancies between the placement of the clade 2 Argentine strains with sequevar reference strains once again highlights the recognized need for systematic and careful modernization of the sub-phylotype taxonomy of RSSC strains [9, 10]. Although the CIP120 assembly (GCF_001644795.1) is fragmented (145 contigs) and slightly less complete (96.2% CheckM completeness) than the genomes analyzed here, it was consistent placed far from other IIA-38 strains in both *egl*-based phylogenetic analyses (Figure 2) and genomic comparisons (Figure 3 and 4). Thus, because genomic analyses and *egl* analysis consistently demonstrated that CIP120 and CFBP6801 are genetically distinct, we propose that sequevar 38 is not a monophyletic lineage. It may be prudent for the next generation of RSSC taxonomy to designate a single strain as “type material,” similar to the approach for species-level taxonomy.

**Figure 2.**
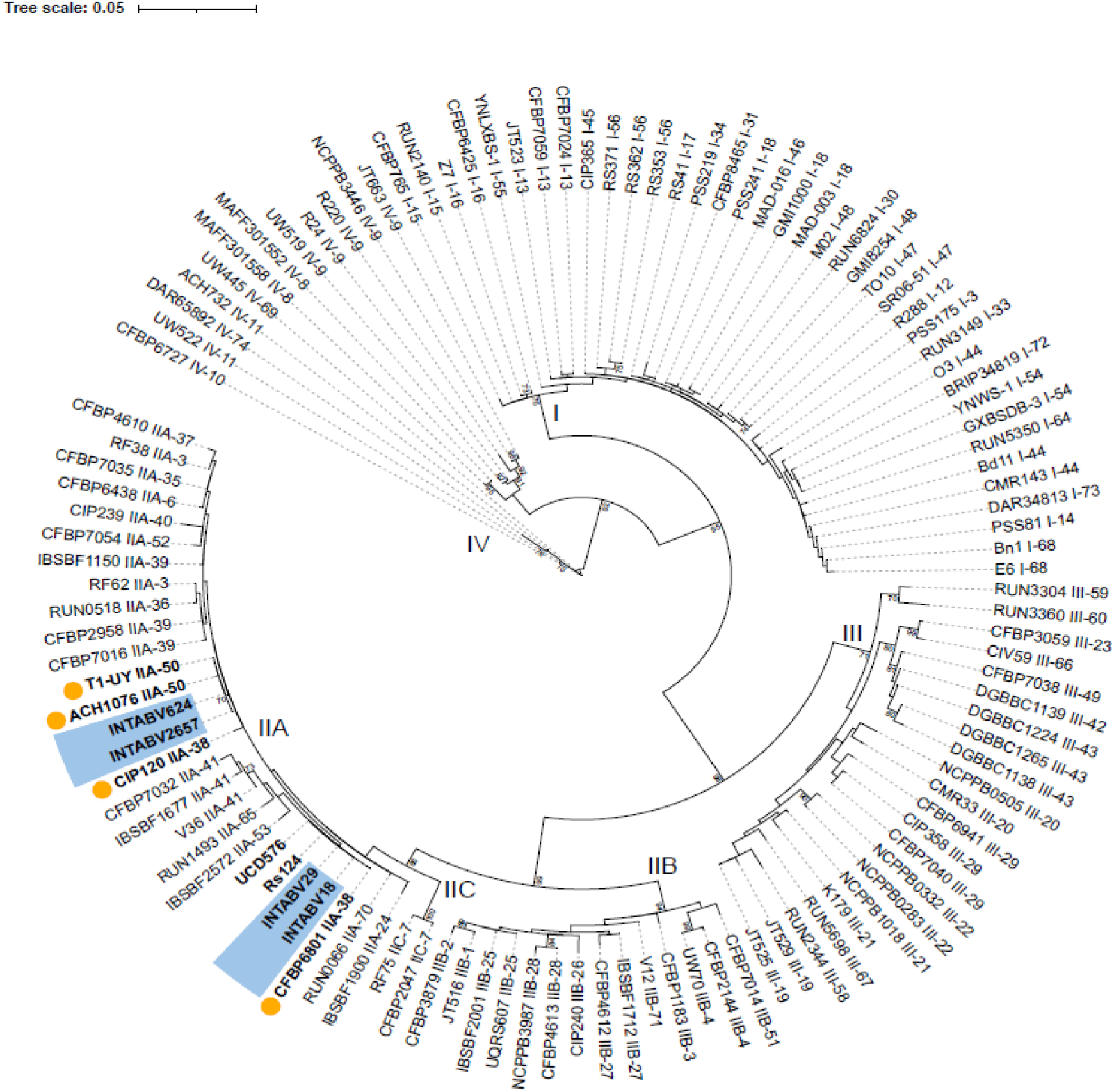
Maximum-likelihood phylogenetic tree inferred from 710 bp of endoglucanase (*egl*) gene sequences assigned Argentine strains as phylotype II sequevar 38 and sequevar 50. The phylogenetic tree was constructed using PhyML v3.0 under the GTR+R nucleotide substitution model, as selected by the SMART model selection procedure (Lefort et al., 2017). *egl* sequences from four Argentine strains (INTABV18, INTABV29, INTABV624, and INTABV2657) are shown in bold and highlighted in blue. Reference *egl* sequences representing sequevars IIA-38 (CFBP6801 and CIP120) and IIA-50 (T1-UY and ACH1076) are also shown in bold and marked with yellow circles. Two USA strains identified as IIA-38 (UCD576 and RS124) are shown in bold. A searchable PDF of this tree in rectangular format is available on Zenodo (doi.org/10.5281/zenodo.19502890).

**Figure 3.**
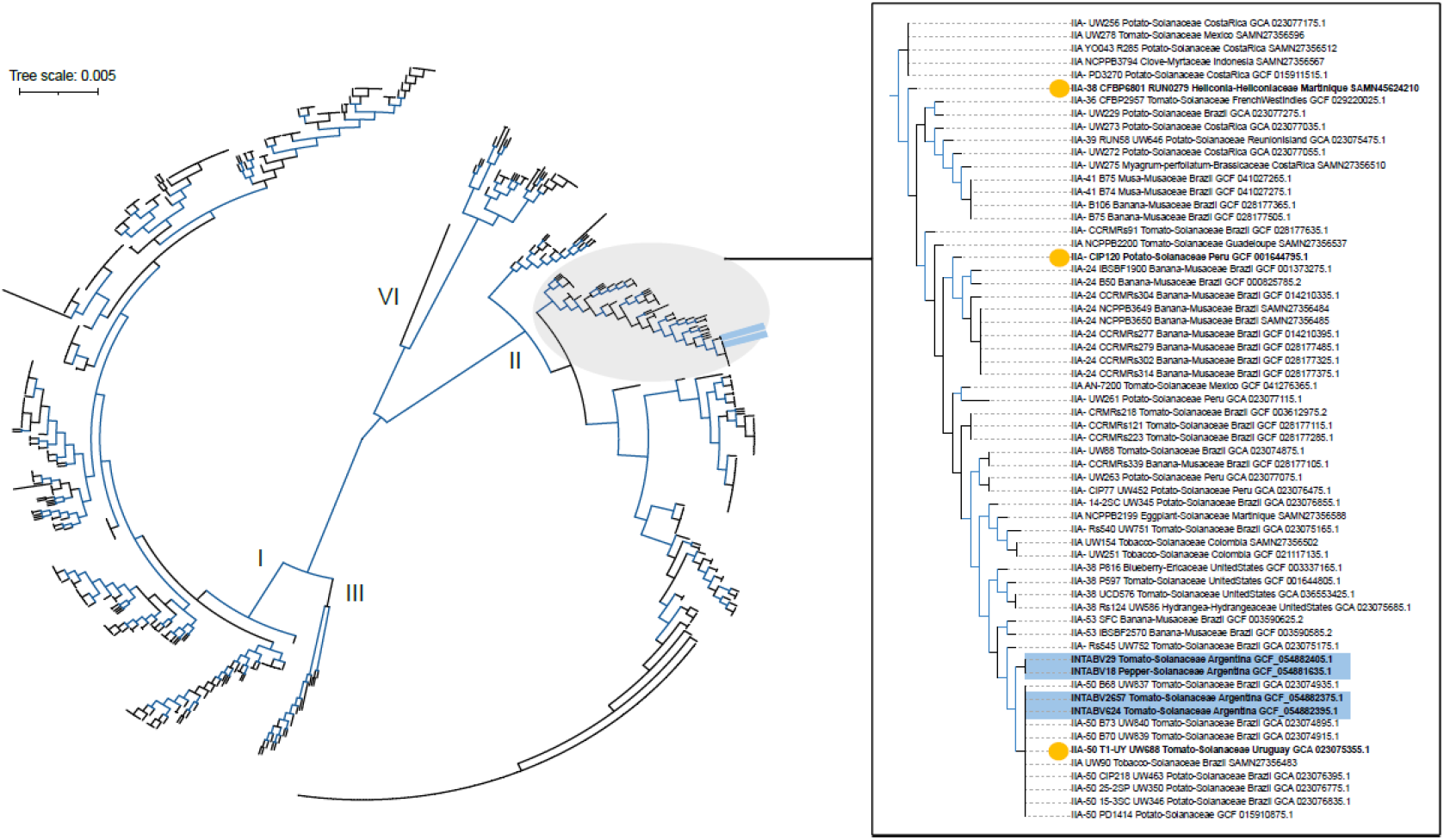
Approximate maximum-likelihood phylogeny based on a concatenated alignment of 49 conserved genes places four Argentine genomes (INTABV18, INTABV29, INTABV624 and INTABV2657) within the phylotype IIA clade. The tree was constructed using the SpeciesTreeBuilder v0.1.4 application on the KBase platform, incorporating the four Argentine genomes into a reference dataset of 825 genomes representing the known global diversity of the RSSC. The tree was visualized and annotated using iTOL v7.4.2. Argentine genomes are shown in bold and highlighted in blue, and *egl* reference strains for the sequevar IIA-38 (CIP120 and CFBP6801) and IIA-50 (T1-UY) are shown in bold and marked with yellow circles. Branches with approximate likelihood-ratio support values higher than >70% are colored in blue. A searchable PDF of this tree in rectangular format is available on Zenodo (doi.org/10.5281/zenodo.19502890).

Although a more systematic epidemiological approach is warranted, we noticed that there were correlations like most Bella Vista isolates were IIA-50 clade (n=6 of 7) and both of the closely related clades were common on tomato. While the 5 eggplant isolates were IIA-50 clade, this may be due to sampling bias as the eggplant isolates were only acquired in Bella Vista where one clade dominated. Importantly, the diversity captured here reflects regional population structure rather than national-scale diversity, as sampling was largely confined to Corrientes province. For example, while the presumed phylotype IIB-1 lineage causing the potato brown rot pandemic has been reported in other Argentine regions [23], this lineage was not present in this dataset. Broader sampling across additional provinces, hosts, and production systems will be required to determine the full distribution of these lineages and the overall diversity of the RSSC in Argentina.

**Figure 4.**
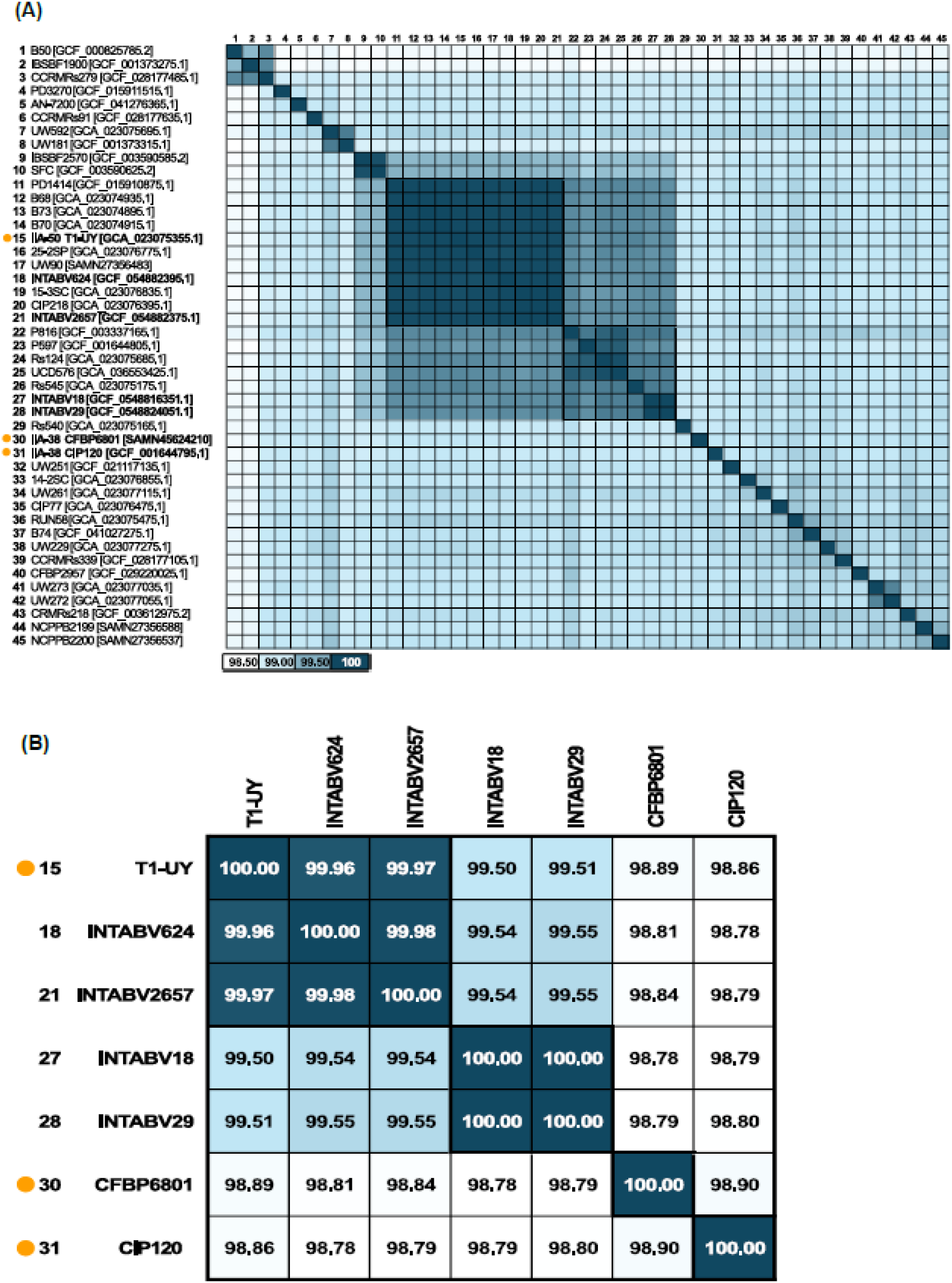
Pairwise Average Nucleotide Identity (ANI) comparisons between the four Argentine genomes and (A) 41 publicly available RSSC genomes and (B) three *egl* reference strains representing sequevars IIA-38 (CIP120 and CFBP6801) and IIA-50 (T1-UY). Reference strains are shown in bold and indicated with yellow circles. INTABV624 and INTABV2657 shared >99.96% ANI with T1-UY. In contrast, INTABV18 and INTABV29 showed ∼98.78–98.80% ANI to the IIA-38 reference strains, but higher similarity to each other (99.54– 99.55%) than to either reference genome. Pairwise ANI values were calculated using fastANI v1.3.3.

## Supporting information

Figure 2_rectangular tree

Figure 3_rectangular tree

Figure S1_rectangular tree

## Conflicts of interest

Authors declare that there are no conflicts of interests

## Funding information

The work was supported in part by INTA-PE.I053 - Generación y difusión de tecnologías para el desarrollo sostenible de los sistemas productivos hortícolas del NEA, the USDA Hatch Program (Project #1023861 to TLP) and the USDA NIFA Pests and Beneficial Species in Agricultural Production Systems (A1112) program (award # 2024-67013-42781 to TLP).

## List of Figures

Figure 1. Map of locations where *Ralstonia solanacearum* species complex has been documented in this study and in the literature.

Figure 2. Maximum-likelihood phylogenetic tree inferred from 710 bp of endoglucanase (*egl*) gene sequences assigned Argentine strains as phylotype II sequevar 38 and sequevar 50.

Figure 3. Approximate maximum-likelihood phylogeny based on a concatenated alignment of 49 conserved genes places four Argentine genomes (INTABV18, INTABV29, INTABV624 and INTABV2657) within the phylotype IIA clade.

Figure 4. Pairwise Average Nucleotide Identity (ANI) comparisons between the four Argentine genomes and (A) 41 publicly available RSSC genomes and (B) three *egl* reference strains representing sequevars IIA-38 (CIP120 and CFBP6801) and IIA-50 (T1-UY).

Figure S1. Maximum-likelihood phylogenetic tree inferred from 471 bp of endoglucanase (*egl*) gene sequences assigned Argentine strains as phylotype II sequevar 38 and sequevar 50.

## List of Tables

Supplementary Table 1. Genomes used in FastANI analysis

Supplementary Table 2. Nanoplot analyses of the raw reads

Supplementary Table 3. QUAST analysis

Supplementary Table 4. CheckM analysis

## Notes

https://doi.org/10.5281/zenodo.19502890

### Competing Interest Statement

The authors have declared no competing interest.

