## Supplementary figures and images for "Phylogenomic Taxonomic Analysis of *Ralstonia solanacearum* Strains causing Bacterial Wilt Disease in Northeastern Argentina"

### Figure 2_rectangular tree

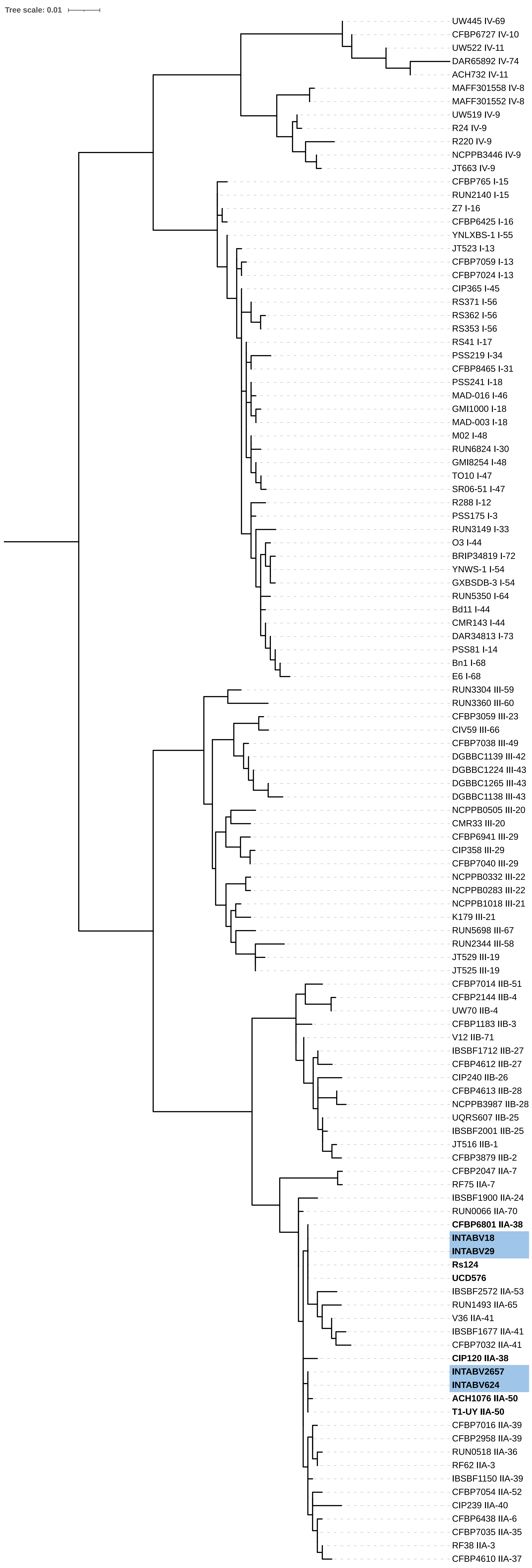

### Figure 3_rectangular tree

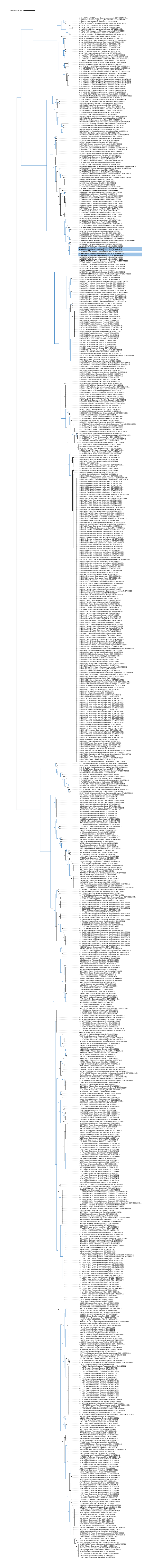

### Figure S1_rectangular tree

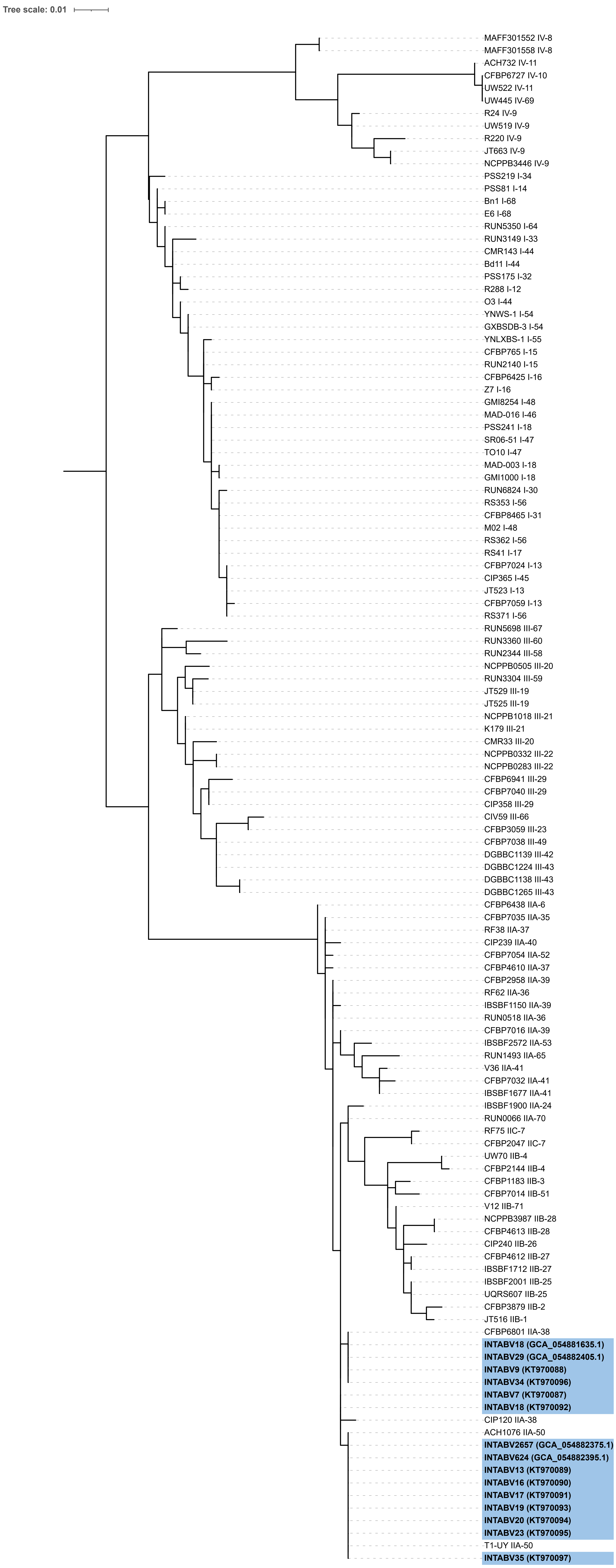
